# Best Practice Data Life Cycle Approaches for the Life Sciences

**DOI:** 10.1101/167619

**Authors:** Philippa C. Griffin, Jyoti Khadake, Kate S. LeMay, Suzanna E. Lewis, Sandra Orchard, Andrew Pask, Bernard Pope, Ute Roessner, Keith Russell, Torsten Seemann, Andrew Treloar, Sonika Tyagi, Jeffrey H. Christiansen, Saravanan Dayalan, Simon Gladman, Sandra B. Hangartner, Helen L. Hayden, William W. H. Ho, Gabriel Keeble-Gagnère, Pasi K. Korhonen, Peter Neish, Priscilla R. Prestes, Mark F. Richardson, Nathan S. Watson-Haigh, Kelly L. Wyres, Neil D. Young, Maria Victoria Schneider

## Abstract

Throughout history, the life sciences have been revolutionised by technological advances; in our era this is manifested by advances in instrumentation for data generation, and consequently researchers now routinely handle large amounts of heterogeneous data in digital formats. The simultaneous transitions towards biology as a data science and towards a ‘life cycle’ view of research data pose new challenges. Researchers face a bewildering landscape of data management requirements, recommendations and regulations, without necessarily being able to access data management training or possessing a clear understanding of practical approaches that can assist in data management in their particular research domain.

Here we provide an overview of best practice data life cycle approaches for researchers in the life sciences/bioinformatics space with a particular focus on ‘omics’ datasets and computer-based data processing and analysis. We discuss the different stages of the data life cycle and provide practical suggestions for useful tools and resources to improve data management practices.

## Introduction

Technological data production capacity is revolutionising biology [1] but is not necessarily correlated with the ability to efficiently analyse and integrate data, or with enabling long-term data sharing and reuse. There are selfish as well as altruistic benefits to making research data reusable [2]: it allows one to find and reuse one’s own previously-generated data easily; it is associated with higher citation rates [3,4]; and it ensures eligibility for funding from and publication in venues that mandate data sharing, an increasingly common requirement [e.g. 5,6,7]. Currently we are losing data at a rapid rate, with up to 80% unavailable after 20 years [8]. This affects reproducibility - assessing the robustness of scientific conclusions by ensuring experiments and findings can be reproduced - which underpins the scientific method. Once access to the underlying data is lost, replicability, reproducibility and extensibility [9] are reduced.

At a broader societal level, the full value of research data may go beyond the initial use case in unforeseen ways [10,11], so ensuring data quality and reusability is crucial to realising its potential value [12–15]. The recent publication of the FAIR principles [12,16] identifies four key criteria for high-quality research data: the data should be Findable, Accessible, Interoperable and Reusable. Whereas a traditional view of data focuses on collecting, processing, analysing data and publishing results only, a life cycle view reveals the additional importance of finding, storing and sharing data [14]. Throughout this article we present a researcher-focused data life cycle framework that has commonalities with other published frameworks [14,17–20] but is aimed at life science researchers specifically (Fig. 1).

**Figure 1:**
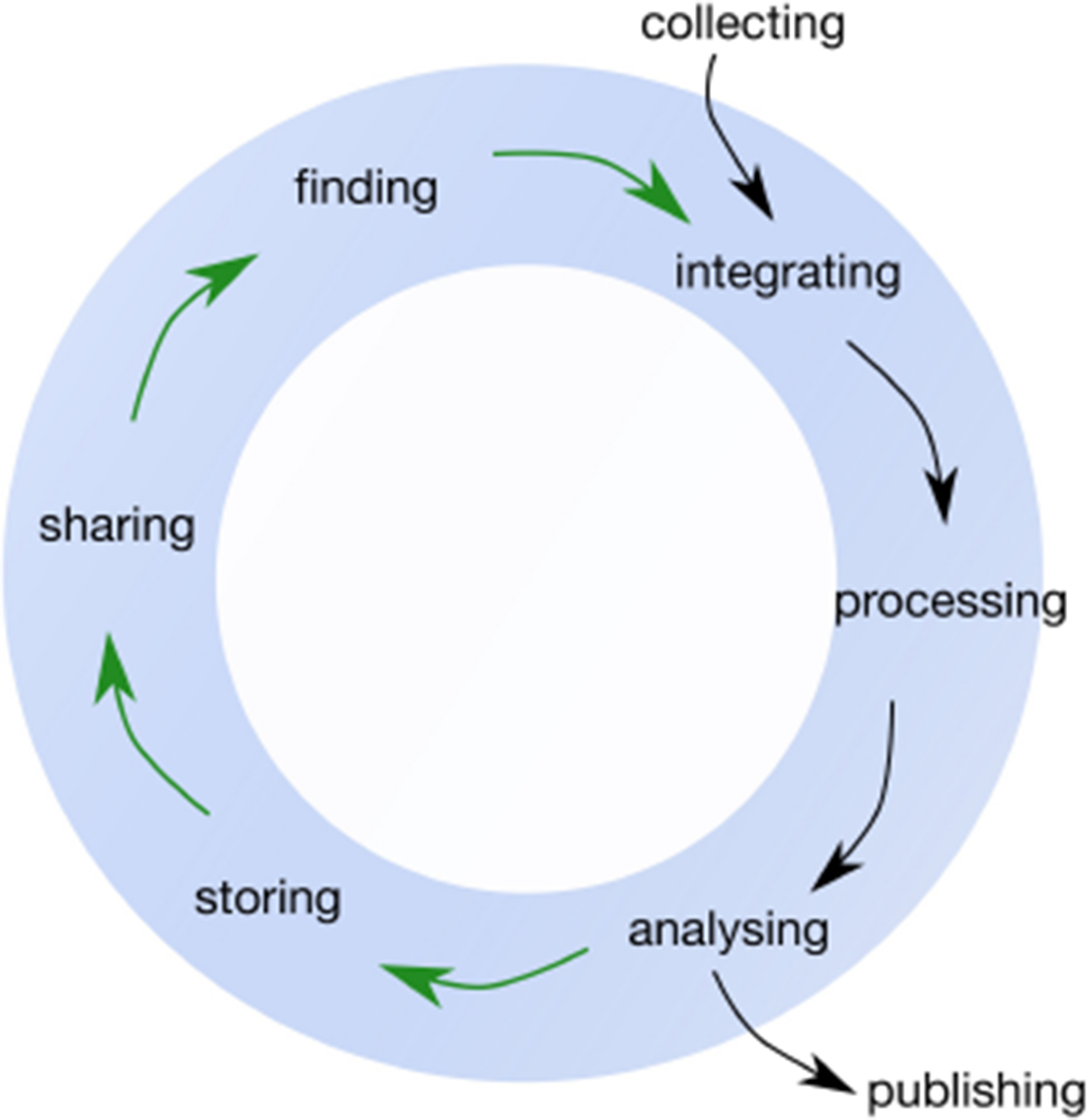
The Data Life Cycle framework for bioscience, biomedical and bioinformatics data that is discussed throughout this article. Black arrows indicate the ‘traditional’, linear view of research data; the green arrows show the steps necessary for data reusability. This framework is likely to be a simplified representation of any given research project, and in practice there would be numerous ‘feedback loops’ and revisiting of previous stages. In addition, the publishing stage can occur at several points in the data life cycle.

Learning how to find, store and share research data is not typically an explicit part of undergraduate or postgraduate training in the biological sciences [21–23]. The scope, size and complexity of datasets in many fields has increased dramatically over the last 10-20 years but the knowledge of how to manage this data is currently limited to specific cohorts of ‘information managers’ (e.g. research data managers, research librarians, database curators and IT professionals with expertise in databases and data schemas [23]). In response to institutional and funding requirements around data availability, a number of tools and educational programs have been developed to help researchers create Data Management Plans to address elements of the data lifecycle [24]; however, even when a plan is mandated, there is often a gap between the plan and the actions of the researcher [13].

During the week of 24-28 October 2016, EMBL Australia Bioinformatics Resource (EMBL- ABR) [25] led workshops on the data life cycle for life science researchers working in the plant, animal, microbial and medical domains. The workshops provided opportunities to (i) map the current approaches to the data life cycle in biology and bioinformatics, and (ii) present and discuss best practice approaches and standards for key international projects with Australian life scientists and bioinformaticians. Discussions during these workshops have informed this publication, which targets life science researchers wanting to improve their data management practice; throughout we highlight some specific data management challenges mentioned by participants.

## Finding Data

In biology, research data is frequently published as supplementary material to articles, on personal or institutional websites, or in non-discipline-specific repositories like Figshare [26] and Dryad [27,28]. In such cases, data may exist behind a paywall, there is no guarantee it will remain extant, and, unless one already knows it exists and its exact location, it may remain undiscovered [29]. It is only when a dataset is added to public data repositories, along with accompanying standardized descriptive metadata (see **Collecting Data**), that it can be indexed and made publicly available [30]. Data repositories also provide unique identifiers that increase findability by enabling persistent linking from other locations and permanent association between data and its metadata.

In the field of molecular biology, a number of bioinformatics-relevant organisations host public data repositories. National and international-level organisations of this kind include the European Bioinformatics Institute (EMBL-EBI) [31], the National Centre for Biotechnology Information (NCBI) [32], the DNA Data Bank of Japan (DDBJ) [33], the Swiss Institute of Bioinformatics (SIB) [34], and the four data center members of the worldwide Protein Data Bank [35], which mirror their shared data with regular, frequent updates. This shared central infrastructure is hugely valuable to research and development. For example, EMBL-EBI resources have been valued at over £270 million per year and contribute to ~£1 billion in research efficiencies; a 20-fold return on investment [36].

Numerous repositories are available for biological data (see Table 1 for an overview), though repositories are still lacking for some data types and sub-domains [37]. Many specialised data repositories exist outside of the shared central infrastructure mentioned, often run voluntarily or with minimal funding. Support for biocuration, hosting and maintenance of these smaller-scale but key resources is a pressing problem [38–40]. The quality of the user-submitted data in public repositories [41,42] can mean that public datasets require extra curation before reuse. Unfortunately, due to low uptake of established methods [43–45] to correct the data [42], the results of extra curation may not find their way back into the repositories. Repositories are often not easily searched by generic web search engines [37]. Registries, which form a secondary layer linking multiple, primary repositories, may offer a more convenient way to search across multiple repositories for data relevant to a researcher’s topics of interest [46].

**Table 1:**
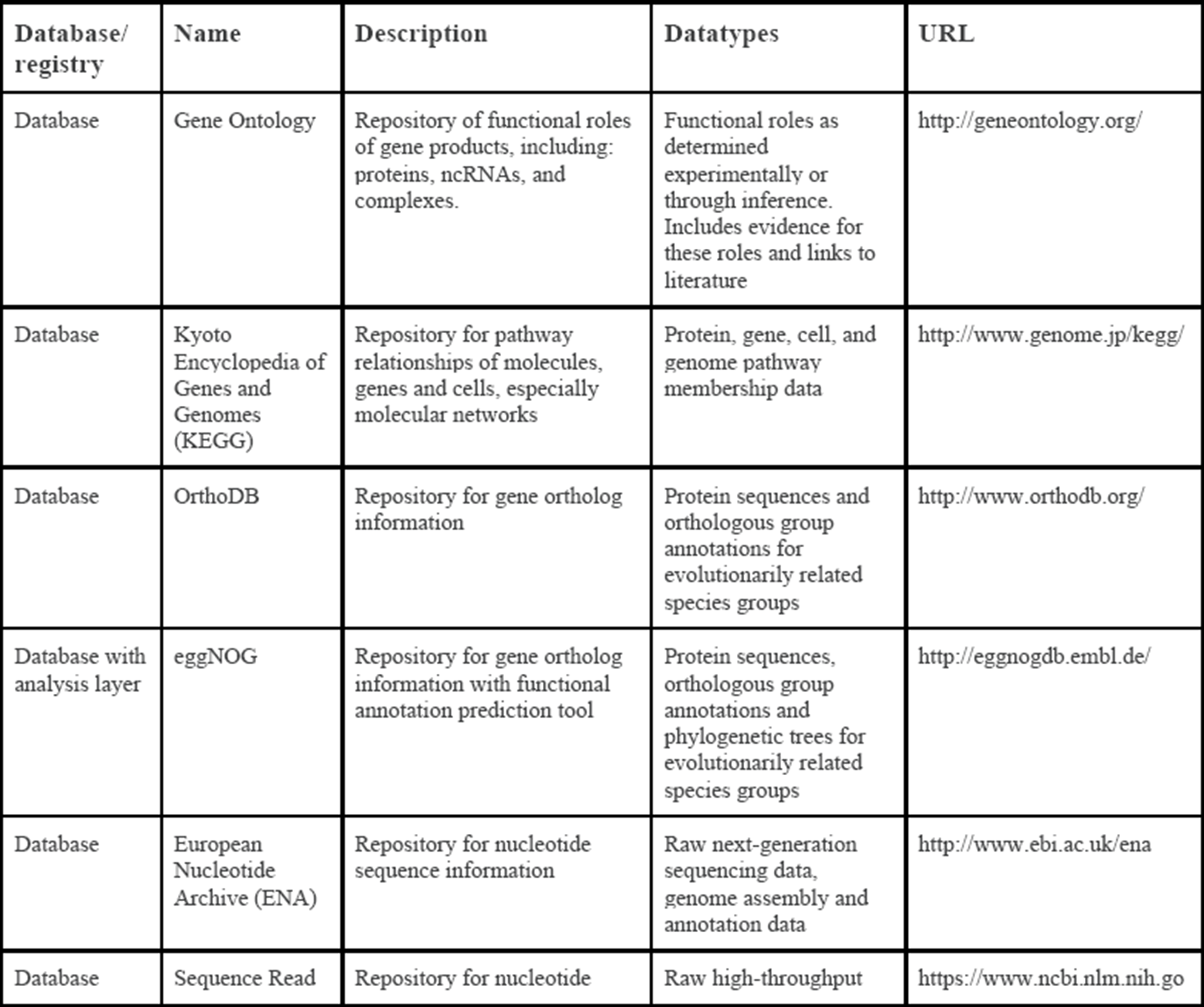
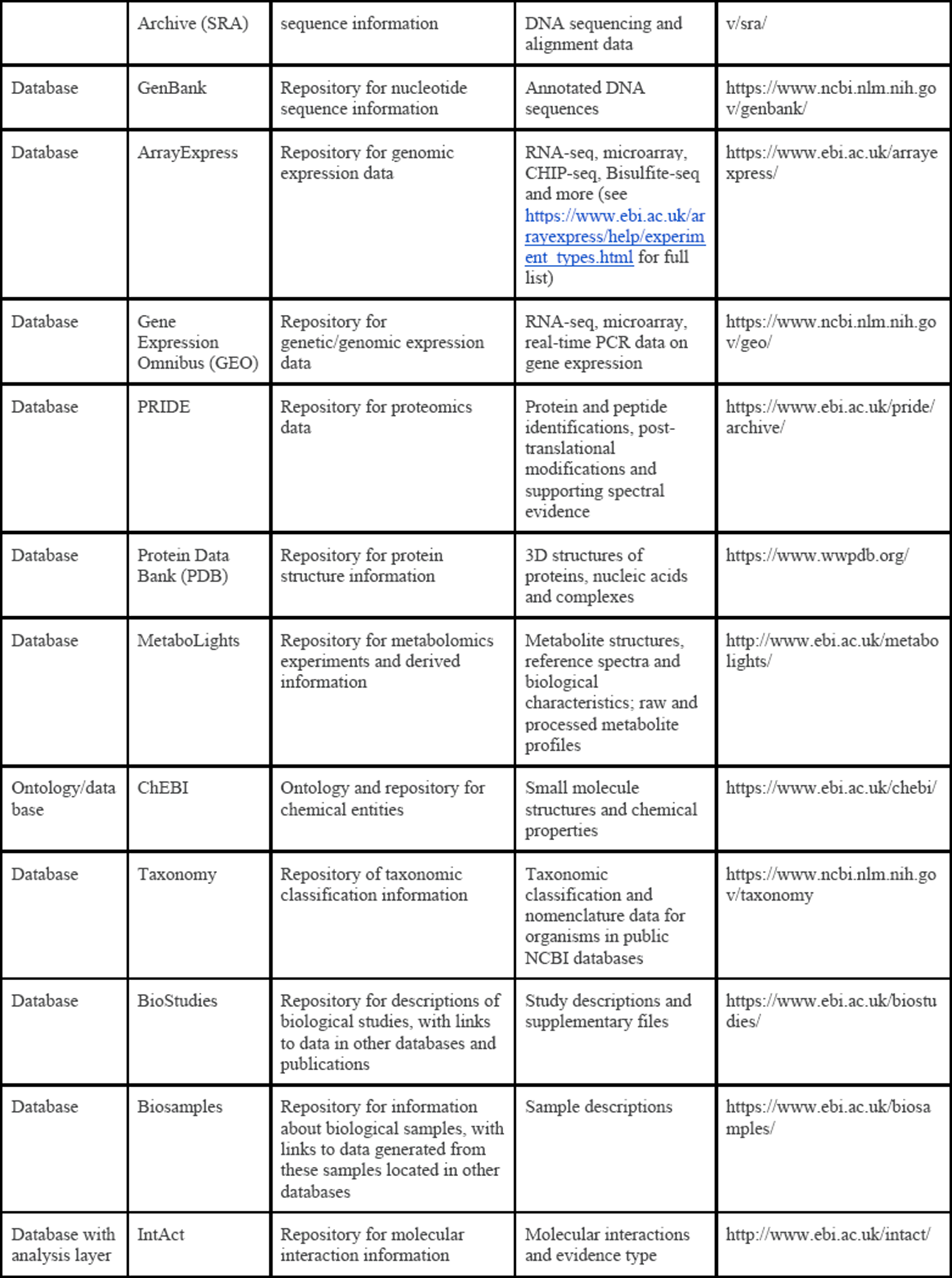
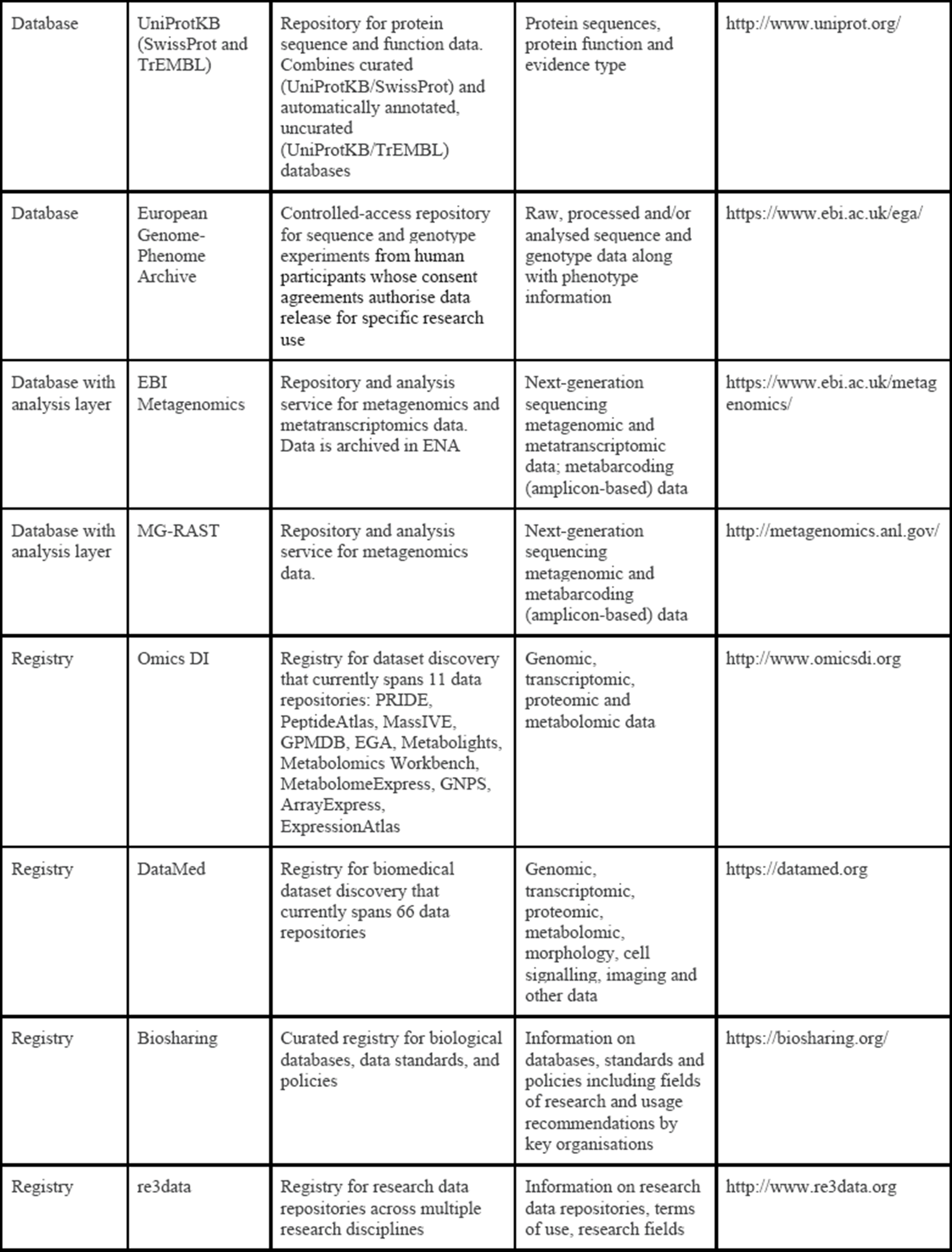
Overview of some representative databases, registries and other tools to find life science data

## Collecting Data

The most useful data has associated information about its creation, its content and its context - called metadata [47]. If metadata is well structured, uses consistent element names and contains element values with specific descriptions from agreed-upon vocabularies, it enables machine readability, aggregation, integration and tracking across datasets: allowing for Findability, Interoperability and Reusability [12,37]. One key approach in best-practice metadata collection is to use controlled vocabularies built from ontology terms. Biological ontologies are tools that provide machine-interpretable representations of some aspect of biological reality [37,48]. They are a way of organising and defining objects (i.e. physical entities or processes), and the relationships between them. Sourcing metadata element values from ontologies ensures that the terms used in metadata are consistent and clearly defined. There are several user-friendly tools available to assist researchers in accessing, using and contributing to ontologies (Table 2).

**Table 2:**
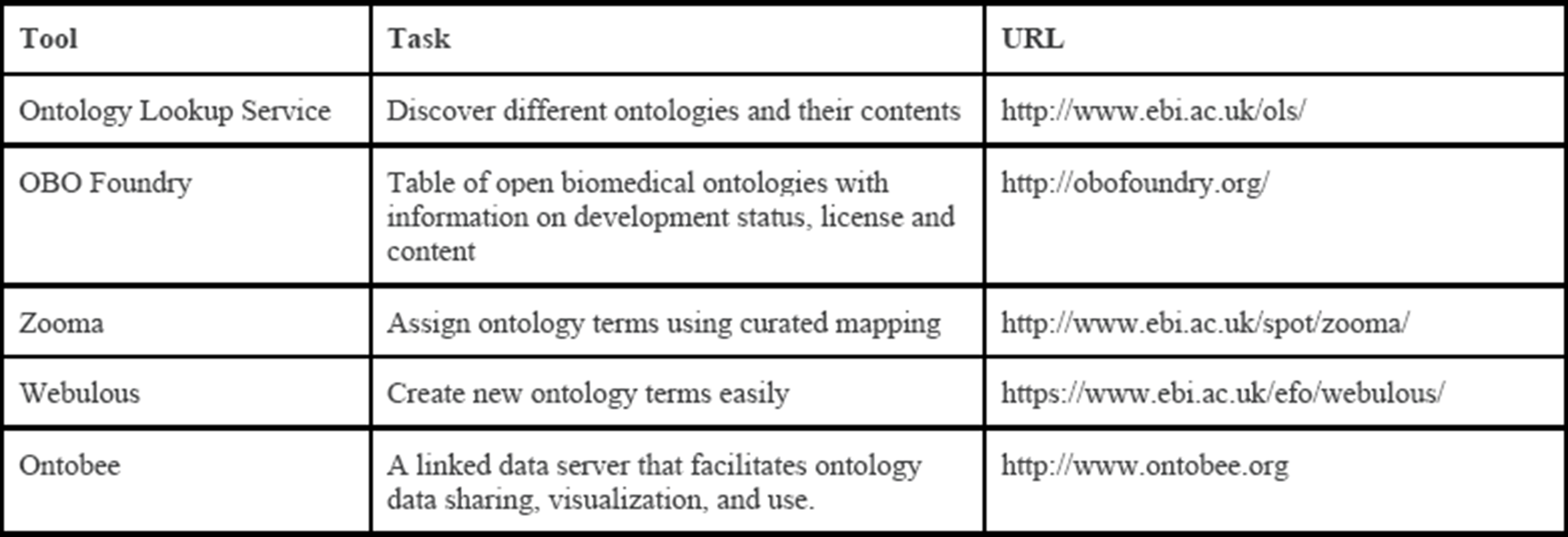
Useful ontology tools to assist in metadata collection

Adopting standard data and metadata formats and syntax is critical for compliance with FAIR principles [12,30,37,46,49]. Biological and biomedical research has been considered an especially challenging research field in this regard, as datatypes are extremely heterogeneous and not all have defined data standards [49,50]; many existing data standards are complex and therefore difficult to use [50], or only informally defined, and therefore subject to variation, misrepresentation, and divergence over time [49]. Nevertheless, well-established standards exist for a variety of biological data types (Table 3). BioSharing [51] is a useful registry of data standards and policies that also indicates the current status of standards for different data types and those recommended by databases and research organisations [46].

**Table 3:**
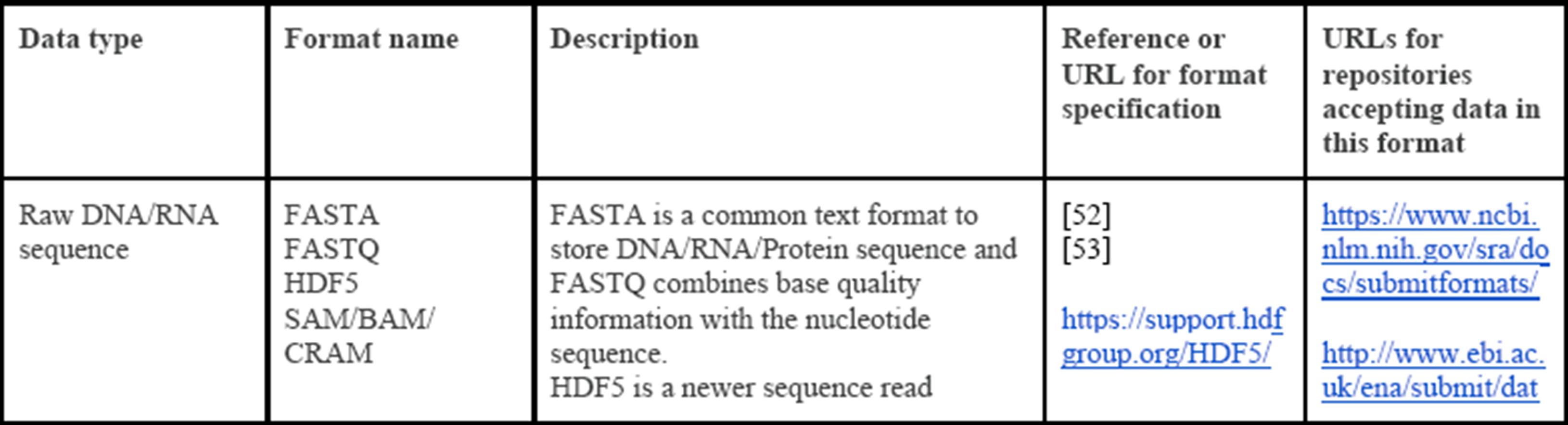
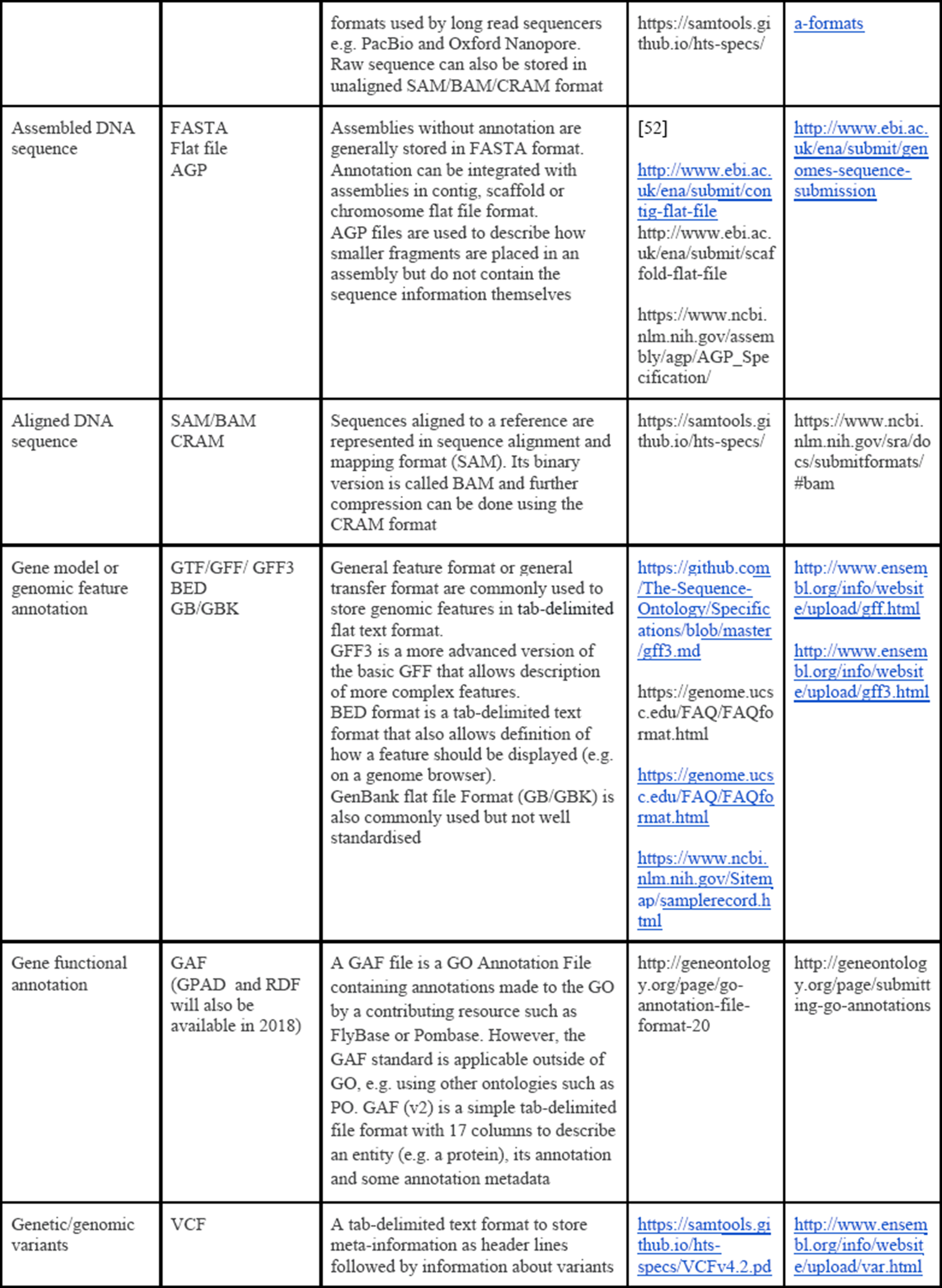
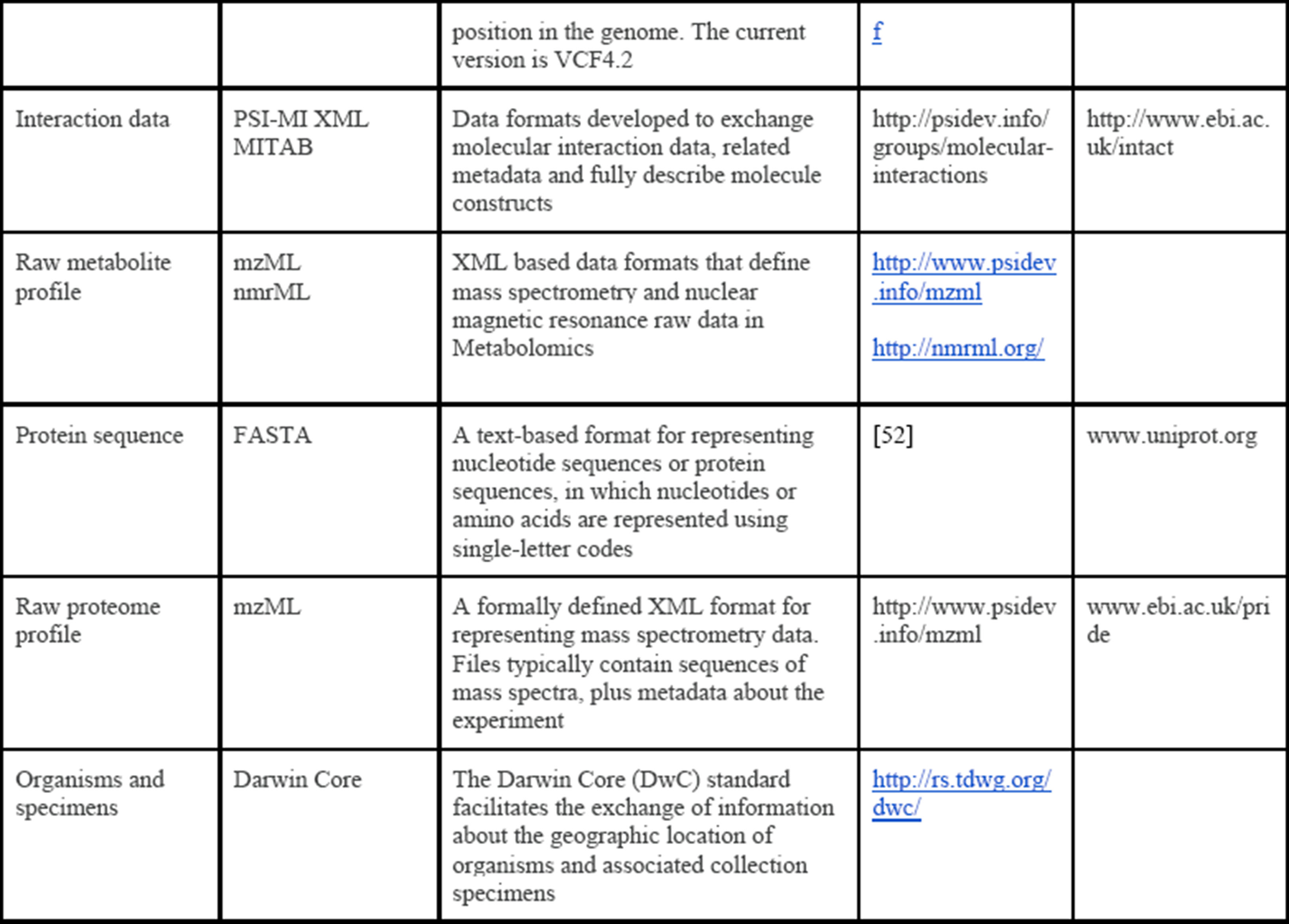
Overview of common standard data formats for ‘omics data

Most public repositories for biological data (see Table 1 and **Storing Data** section) require that minimum metadata be submitted accompanying each dataset (Table 4). This minimum metadata specification typically has broad community input [54]. Minimum metadata standards may not include the crucial metadata fields that give the full context of the particular research project [54], so it is important to gather metadata early, understand how to extend a minimum metadata template to include additional fields in a structured way, and think carefully about all the relevant pieces of metadata information that might be required for reuse.

**Table 4:**
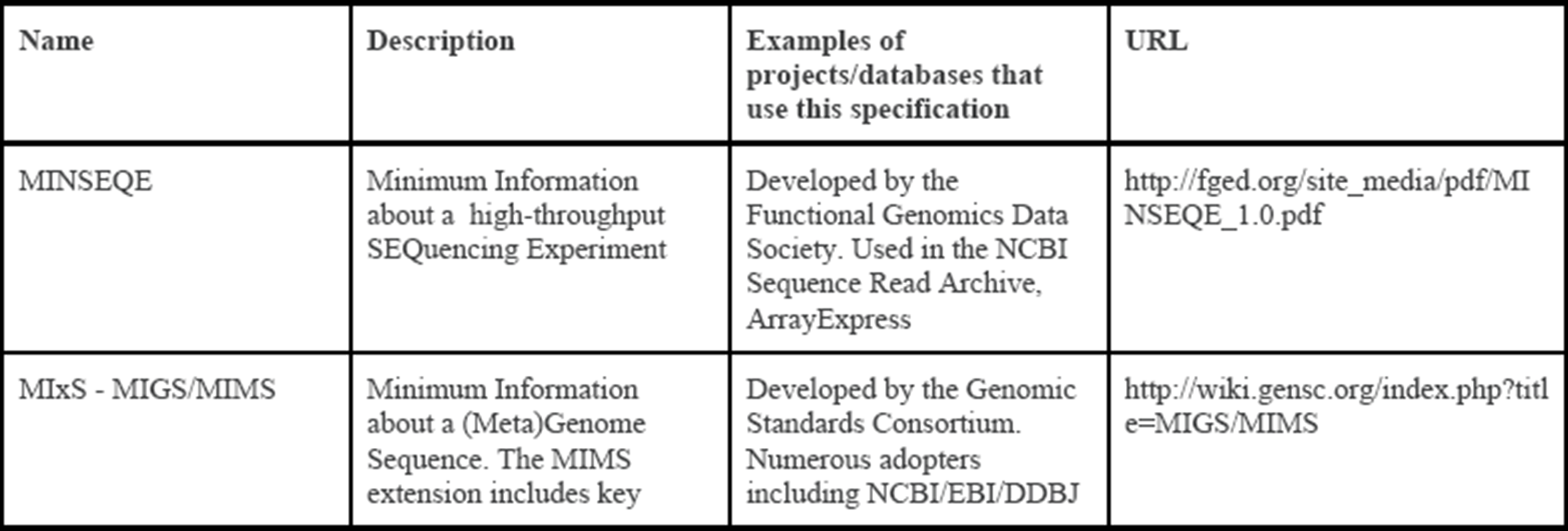
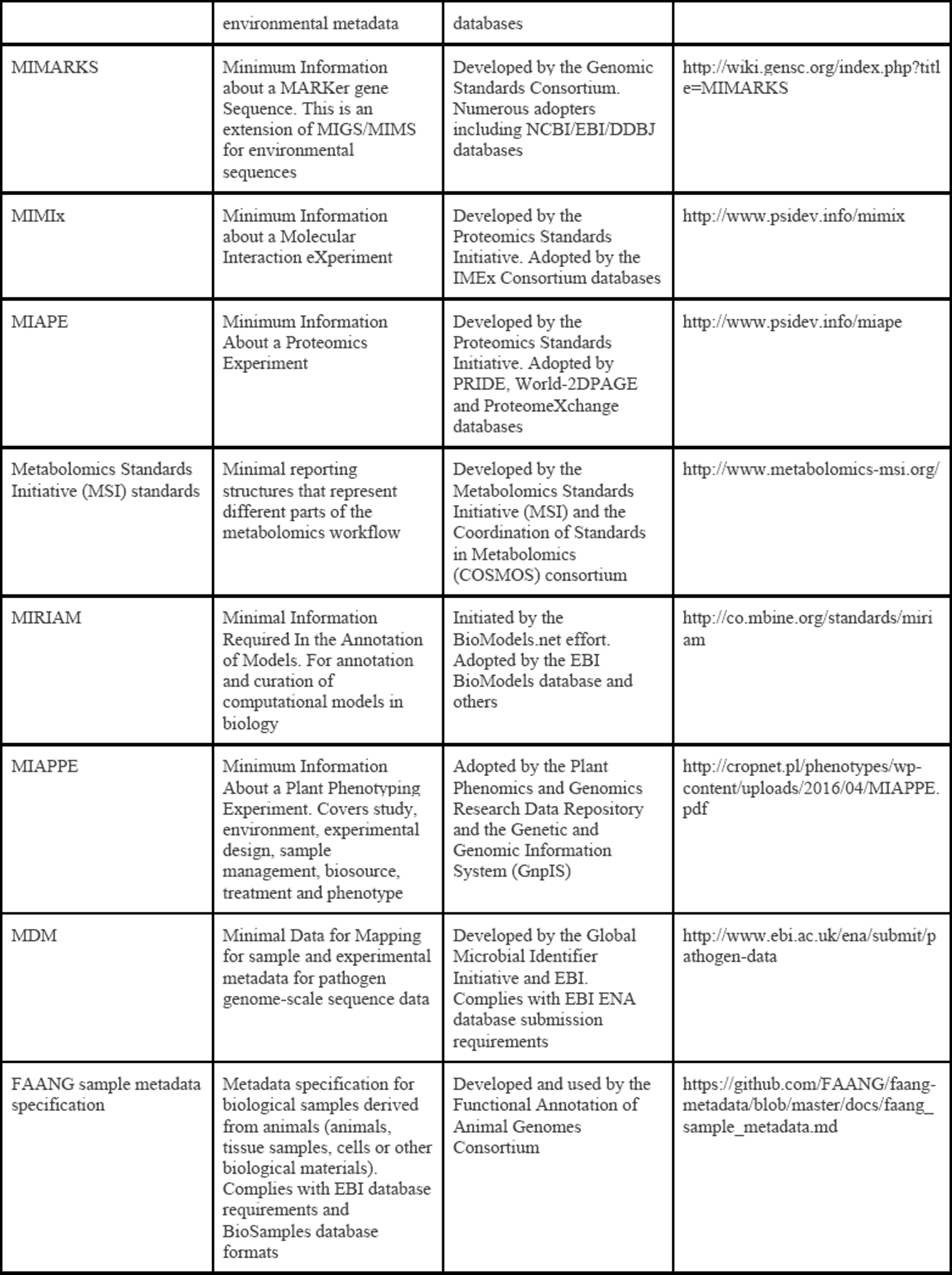
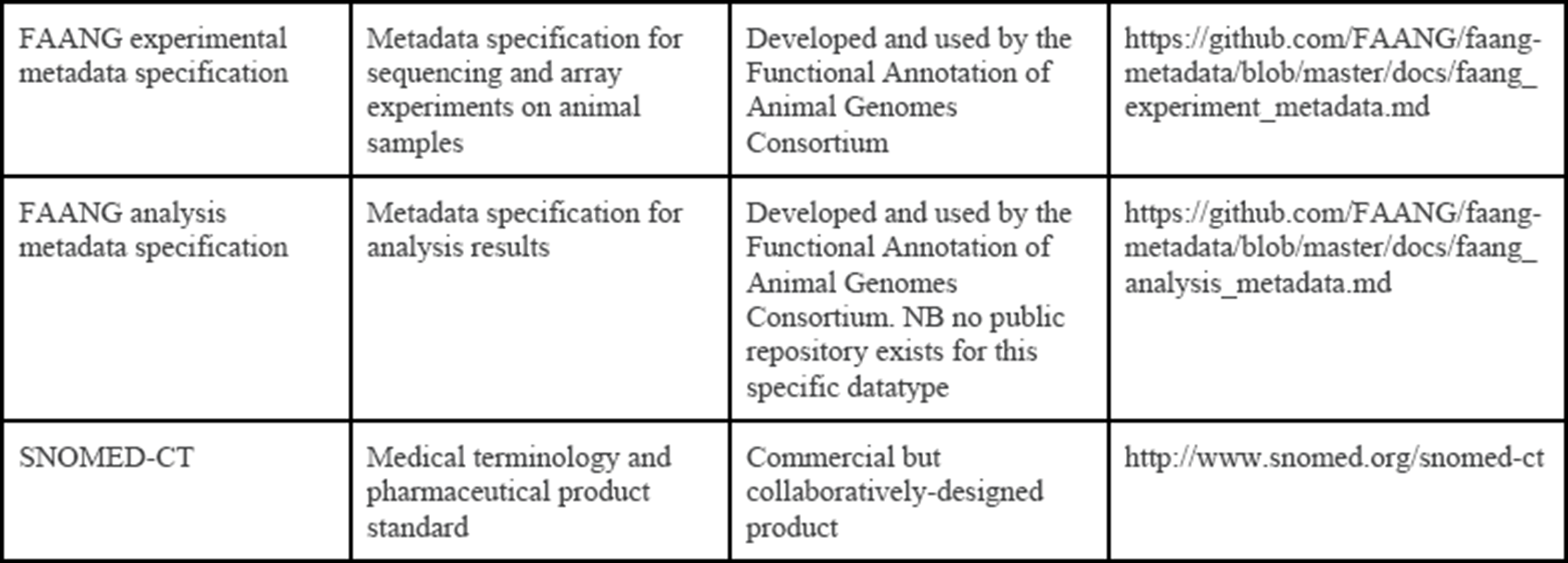
Some community-designed minimum information criteria for metadata specifications in life sciences

## Processing & Analysing Data

Recording and reporting how research data is processed and analysed computationally is crucial for reproducibility and assessment of research quality [1,55]. Full reproducibility requires access to the software, software versions, dependencies and operating system used as well as the data and software code itself [56]. Therefore, although computational work is often seen as enabling reproducibility in the short term, in the long term it is fragile and reproducibility is limited [57–59]. Best-practice approaches for preserving data processing and analysis code involve hosting source code in a repository where it receives a unique identifier and is under version control; where it is open, accessible, interoperable and reusable - broadly mapping to the FAIR principles for data. Github [60] and Bitbucket [61], for example, fulfil these criteria, and Zenodo additionally generates Digital Object Identifiers (DOIs) for submissions and guarantees long-term archiving [62]. Several recent publications have suggested ways to improve current practice in research software development [20,63–65].

The same points hold for wet-lab data production: for full reproducibility, it is important to capture and enable access to specimen cell lines, tissue samples and/or DNA as well as reagents. Wet-lab methods can be captured in electronic laboratory notebooks and reported in the Biosamples database [66], protocols.io [67] or OpenWetWare [68]; specimens can be lodged in biobanks, culture or museum collections [69–73]; but the effort involved in enabling full reproducibility remains extensive. Electronic laboratory notebooks are frequently suggested as a sensible way to make this information openly available and archived [74]. Some partial solutions exist [e.g. 75,76–78], including tools for specific domains such as the Scratchpad Virtual Research Environment for natural history research [79]. Other tools can act as or be combined to produce notebooks for small standalone code-based projects [80,81], including Jupyter Notebook [e.g. 82], Rmarkdown [83], and Docker [84]. However, it remains a challenge to implement online laboratory notebooks to cover both field/lab work and computer-based work, especially when computer work is extensive, involved and non-modular [55]. Currently, no best-practice guidelines or minimum information standards exist for use of electronic laboratory notebooks [9]. We suggest that appropriate minimum information to be recorded for most computer-based tasks should include date, task name and brief description, aim, actual command(s) used, software names and versions used, input/output file names and locations, script names and locations.

During the EMBL-ABR workshop series, participants identified the data processing and analysis stage as one of the most challenging for openness. A few participants had put intensive individual effort into developing custom online lab (and code) notebook approaches but the majority had little awareness of this as a useful goal. This suggests a gap between modern biological research as a field of data science, and biology as it is still mostly taught in undergraduate courses, with little or no focus on computational analysis, or project or data management. As reported elsewhere [21–23], this gap has left researchers lacking key knowledge and skills required to implement best practices in dealing with the life cycle of their data.

## Publishing Data

Traditionally, scientific publications included raw research data, but in recent times datasets have grown beyond the scope of practical inclusion in a manuscript [14,55]. Selected data outputs are often included without sharing or publishing the underlying raw data [19]. Journals increasingly recommend or require deposition of raw data in a public repository [e.g. 85], although exceptions have been made for publications containing commercially-relevant data [86]. The current data-sharing mandate is somewhat field-dependent [8,87] and also varies within fields [88]. For example, in the field of bioinformatics, the UPSIDE principle [89] is referred to by some journals (e.g. Bioinformatics [90]), while others have journal- or publisher-specific policies (e.g. BMC Bioinformatics [91]).

The vast majority of scientific journals require inclusion of processing and analysis methods in ‘sufficient detail for reproduction’ [e.g. 92,93–97], though journal requirements are diverse and complex [98], and the level of detail authors provide can vary greatly in practice [99,100]. More recently, many authors have highlighted that full reproducibility requires sharing data and resources at all stages of the scientific process, from raw data (including biological samples) to full methods and analysis workflows [1,9,72,100]. This remains a challenge however [101,102], as discussed in the **Processing and Analysing Data** section. To our knowledge, strategies for enabling computational reproducibility are currently not mandated by any scientific journal.

A recent development in the field of scientific publishing is the establishment of ‘data journals’: scientific journals that publish papers describing datasets. This gives authors a vehicle to accrue citations (still a dominant metric of academic impact) for data production alone, which can often be labour-intensive and expensive yet is typically not well recognised under the traditional publishing model. Examples of this article type include the Data Descriptor in Scientific Data [103] and the Data Note in GigaScience [104], which do not include detailed new analysis but rather focus on describing and enabling reuse of datasets.

The movement towards sharing research publications themselves (‘Open Access Publishing’) has been discussed extensively elsewhere [e.g. 29,105,106]. Publications have associated metadata [creator, date, title etc.; 107] and unique identifiers (PubMed ID for biomedical and some life science journals, DOIs for the vast majority of journals; see Table 5). The ORCID system [108] enables researchers to claim their own unique identifier, which can be linked to their publications. The use of unique identifiers within publications referring to repository records (e.g. genes, proteins, chemical entities) is not generally mandated by journals [e.g. 109], although it would ensure a common vocabulary is used and so make scientific results more interoperable and reusable [110]. Some efforts are underway to make this easier for researchers: for example, Genetics and other Genetics Society of America journals assist authors in linking gene names to model organism database entries [111].

**Table 5:** Identifiers throughout the data life cycle

| Name | Relevant stage of data life cycle | Description | URL |
| --- | --- | --- | --- |
| Digital Object Identifier (DOI) | Publishing, Sharing, Finding | A unique identifier for a digital (or physical or abstract) object | https://www.doi.org/ |
| Open Researcher and Contributor ID (ORCID) | Publishing | An identifier for a specific researcher that persists across publications and other research outputs | https://orcid.org/ |
| Repository accession number | Finding, Processing/Analyzing, Publishing, Sharing, Storing | A unique identifier for a record within a repository. Format will be repository-specific. Examples include NIH UIDs (unique identifiers) and accession numbers; ENA accession numbers; PDB IDs | For example, https://support.ncbi.nlm.nih.gov/link/portal/28045/28049/Article/499/<br>http://www.ebi.ac.uk/ena/submit/accession-number-formats |
| Pubmed ID (PMID) | Publishing | An example of a repository-specific unique identifier: PubMed IDs are used for research publications indexed in the PubMed database | https://www.ncbi.nlm.nih.gov/pubmed/ |
| International Standard Serial Number (ISSN) | Publishing | A unique identifier for a journal, magazine or periodical | http://www.issn.org/ |
| International Standard Book Number (ISBN) | Publishing | A unique identifier for a book, specific to the title, edition and format | https://www.isbn-international.org |

## Storing Data

While primary data archives are the best location for raw data and some downstream data outputs (Table 1), researchers also need local data storage solutions during the processing and analysis stages. Data storage requirements vary among research domains, with major challenges often evident for groups working on taxa with large genomes (e.g. crop plants), which require large storage resources, or on human data, where privacy regulations may require local data storage, access controls and conversion to non-identifiable data if data is to be shared [112–114]. In addition, long-term preservation of research data should consider threats such as storage failure, mistaken erasure, bit rot, outdated media, outdated formats, loss of context and organisational failure [115].

## Sharing Data

The best-practice approach to sharing biological data is to deposit it (with associated metadata) in a primary archive suitable for that datatype [11] that complies with FAIR principles. As highlighted in the **Storing Data** section, these archives assure both data storage and public sharing as their core mission, making them the most reliable location for long-term data storage. Alternative data sharing venues (e.g. FigShare, Dryad) do not require or implement specific metadata or data standards. This means that while these venues have a low barrier to entry for submitters, the data is not FAIR unless submitters have independently decided to comply with more stringent criteria. If available, an institutional repository may be a good option if there is no suitable archive for that datatype. Importantly, plans for data sharing should be made at the start of a research project and reviewed during the project, to ensure ethical approval is in place and that the resources and metadata needed for effective sharing are available at earlier stages of the data life cycle [3].

During the EMBL-ABR workshop series, the majority of participants were familiar with at least some public primary data repositories, and many had submitted data to them previously. A common complaint was around usability of current data submission tools and a lack of transparency around metadata requirements and the rationale for them. A few workshop participants raised specific issues about the potential limitations of public data repositories where their data departed from the assumptions of the repository (e.g. unusual gene models supported by experimental evidence that were rejected by the automated NCBI curation system). Most workshop participants were unaware they could provide feedback to the repositories to deal with such situations, and this could also be made clearer on the repository websites. Again, this points in part to existing limitations in the undergraduate and postgraduate training received by researchers, where the concepts presented in this article are presented as afterthoughts, if at all. On the repository side, while there is a lot of useful information and training material available to guide researchers through the submission process [e.g. the EMBL-EBI Train Online webinars and online training modules, 116], it is not always linked clearly from the database portals or submission pages themselves. Similarly, while there are specifications and standards available for many kinds of metadata [Table 4; also see 51], many do not have example templates available, which would assist researchers in implementing the standards in practice.

### What can the research community do to encourage best-practice?

We believe that the biological/biomedical community and individual researchers have a responsibility to the public to help advance knowledge by making research data FAIR for reuse [12], especially if the data were generated using public funding. There are several steps that can assist in this mission:

1. **Senior scientists should lead by example** and ensure all the data generated by their laboratories is well-managed, fully annotated with the appropriate metadata and made publicly available in an appropriate repository.
2. **The importance of data management and benefits of data reuse should be taught** at the undergraduate and postgraduate levels [23]. Computational biology and bioinformatics courses in particular should include material about data repositories, data and metadata standards, data discovery and access strategies. Material should be domain-specific enough for students to attain learning outcomes directly relevant to their research field.
3. Funding bodies are already taking a lead role in this area by requiring the incorporation of a data management plan into grant applications. A next step would be for a **formal check, at the end of the grant period, that this plan has been adhered to and data is available in an appropriate format for reuse** [13].
4. **Funding bodies and research institutions should judge quality dataset generation as a valued metric when evaluating grant or promotion applications**.
5. **Similarly, leadership and participation in community efforts in data and metadata standards, and open software and workflow development should be recognised as academic outputs**.
6. **Data repositories should ensure that the data deposition and third-party annotation processes are as FAIR and painless as possible** to the naive researcher, without the need for extensive bioinformatics support [42].
7. **Journals should require editors and reviewers to check manuscripts to ensure that all data, including research software code and samples where appropriate, have been made publicly available in an appropriate repository**, and that methods have been described in enough detail to allow re-use and meaningful reanalysis [11].
8. Finally, **researchers reusing any data should openly acknowledge this fact and fully cite the dataset, including unique identifiers** [11,13,37].

## Conclusions

While the concept of a life cycle for research data is appealing from an Open Science perspective, challenges remain for life science researchers to put this into practice. During the EMBL-ABR Data Life Cycle workshop series, we noted limited awareness among attendees of the resources available to researchers that assist in finding, collecting, processing, analysis, publishing, storing and sharing FAIR data. We believe this manuscript provides a useful overview of the relevant concepts and an introduction to key organisations, resources and guidelines to help researchers improve their data management practices.

Furthermore, we note that data management in the era of biology as a data science is a complex and evolving topic and both best practices and challenges are highly domain-specific, even within the life sciences. This factor may not always be appreciated at the organisational level, but has major practical implications for the quality and interoperability of shared life science data. Finally, domain-specific education and training in data management would be of great value to the life science research workforce, and we note an existing gap at the undergraduate, postgraduate and short course level in this area.

## Acknowledgements

The authors thank Dan Bolser for his involvement in the EMBL-ABR Data Life Cycle workshops, and all workshop participants for sharing their experiences and useful discussions.

## References

1. Cohen-Boulakia S, Belhajjame K, Collin O, Chopard J, Froidevaux C, Gaignard A, et al. Scientific workflows for computational reproducibility in the life sciences: status, challenges and opportunities. Future Gener Comput Syst. doi:10.1016/j.future.2017.01.012

2. Hampton SE, Anderson SS, Bagby SC, Gries C, Han X, Hart EM, et al. The Tao of open science for ecology. Ecosphere. 2015;6: 1–13. doi:10.1890/ES14-00402.1

3. Lord P, Macdonald A, Sinnot R, Ecklund D, Westhead M, Jones A. Large-scale data sharing in the life sciences: Data standards, incentives, barriers and funding models [Internet]. UK e-Science; 2005. Report No.: UKeS-2006-02. Available: http://www.nesc.ac.uk/technical_papers/UKeS-2006-02.pdf

4. Piwowar HA, Vision TJ. Data reuse and the open data citation advantage. PeerJ. 2013;1: e175. doi:10.7717/peerj.175

5. National Institutes of Health. NIH Guide: Final NIH statement on sharing research data [Internet]. 2003 [cited 19 May 2017]. Available: https://grants.nih.gov/grants/guide/notice-files/NOT-OD-03-032.html

6. Wellcome Trust. Policy on data management and sharing. In: Wellcome Trust [Internet]. 2017 [cited 19 May 2017]. Available: https://wellcome.ac.uk/funding/managing-grant/policy-data-management-and-sharing

7. Bill & Melinda Gates Foundation. Open access policy. In: Bill & Melinda Gates Foundation [Internet]. 2015 [cited 19 May 2017]. Available: http://www.gatesfoundation.org/How-We-Work/General-Information/Open-Access-Policy

8. Vines TH, Albert AYK, Andrew RL, Débarre F, Bock DG, Franklin MT, et al. The availability of research data declines rapidly with article age. Curr Biol. 2014;24: 94–97. doi:10.1016/j.cub.2013.11.014

9. Lewis J, Breeze CE, Charlesworth J, Maclaren OJ, Cooper J. Where next for the reproducibility agenda in computational biology? BMC Syst Biol. 2016;10: 52. doi:10.1186/s12918-016-0288-x

10. Voytek B. The virtuous cycle of a data ecosystem. PLoS Comput Biol. 2016;12: e1005037. doi:10.1371/journal.pcbi.1005037

11. Whitlock MC. Data archiving in ecology and evolution: best practices. Trends Ecol Evol. 2011;26: 61–65. doi:10.1016/j.tree.2010.11.006

12. Wilkinson MD, Dumontier M, Aalbersberg IJ, Appleton G, Axton M, Baak A, et al. The FAIR Guiding Principles for scientific data management and stewardship. Scientific Data. 2016;3: 160018. doi:10.1038/sdata.2016.18

13. Van Tuyl S, Whitmire AL. Water, water, everywhere: defining and assessing data sharing in academia. PLoS One. 2016;11: e0147942. doi:10.1371/journal.pone.0147942

14. Rüegg J, Gries C, Bond-Lamberty B, Bowen GJ, Felzer BS, McIntyre NE, et al. Completing the data life cycle: using information management in macrosystems ecology research. Front Ecol Environ. Ecological Society of America; 2014;12: 24–30. doi:10.1890/120375

15. Moody D, Walsh P. Measuring the value of information: an asset valuation approach. European Conference on Information Systems. 1999. p. 17.

16. Mons B, Neylon C, Velterop J, Dumontier M, da Silva Santos LOB, Wilkinson MD. Cloudy, increasingly FAIR; revisiting the FAIR Data guiding principles for the European Open Science Cloud. Inf Serv Use. IOS Press; 1–8. doi:10.3233/ISU-170824

17. DataONE. Data Life Cycle. In: DataONE [Internet]. [cited 19 May 2017]. Available: https://www.dataone.org/data-life-cycle

18. Faundeen JL, Burley TE, Carlino JA, Govoni DL, Henkel HS, Holl SL, et al. The United States geological survey science data lifecycle model [Internet]. US Geological Survey; 2014. Available: https://pubs.er.usgs.gov/publication/ofr20131265

19. Michener WK, Jones MB. Ecoinformatics: supporting ecology as a data-intensive science. Trends Ecol Evol. 2012;27: 85–93. doi:10.1016/j.tree.2011.11.016

20. Lenhardt WC, Ahalt S, Blanton B, Christopherson L, Idaszak R. Data management lifecycle and software lifecycle management in the context of conducting science. Journal of Open Research Software. 2014;2: e15. doi:10.5334/jors.ax

21. Campbell. Data’s shameful neglect. Nature. 2009;461: 145. doi:10.1038/461145a

22. Strasser CA, Hampton SE. The fractured lab notebook: undergraduates and ecological data management training in the United States. Ecosphere. Ecological Society of America; 2012;3: 1–18. doi:10.1890/ES12-00139.1

23. Tenopir C, Allard S, Sinha P, Pollock D, Newman J, Dalton E, et al. Data management education from the perspective of scientific educators. International Journal of Digital Curation. 2016;11: 232–251. doi:10.2218/ijdc.v11i.389

24. Simms S, Strong M, Jones S, Ribeiro M. The future of data management planning: tools, policies, and players. International Journal of Digital Curation. 2016;11: 208–217. doi:10.2218/ijdc.v11i1.413

25. Schneider MV, Griffin PC, Tyagi S, Flannery M, Dayalan S, Gladman S, et al. Establishing a distributed national research infrastructure providing bioinformatics support to life science researchers in Australia. Brief Bioinform. 2017; doi:10.1093/bib/bbx071

26. Figshare. In: Figshare [Internet]. 2017 [cited 26 May 2017]. Available: https://figshare.com/

27. Dryad. In: Dryad [Internet]. 2017 [cited 26 May 2017]. Available: http://datadryad.org/

28. Womack RP. Research data in core journals in biology, chemistry, mathematics, and physics. PLoS One. 2015;10: e0143460. doi:10.1371/journal.pone.0143460

29. McKiernan EC, Bourne PE, Brown CT, Buck S, Kenall A, Lin J, et al. How open science helps researchers succeed. Elife 2016;5: e16800. doi:10.7554/eLife.16800

30. Sansone S-A, Rocca-Serra P, Field D, Maguire E, Taylor C, Hofmann O, et al. Toward interoperable bioscience data. Nat Genet. 2012;44: 121–126. doi:10.1038/ng.1054

31. Cook CE, Bergman MT, Finn RD, Cochrane G, Birney E, Apweiler R. The European Bioinformatics Institute in 2016: data growth and integration. Nucleic Acids Res. 2016;44: D20–6. doi:10.1093/nar/gkv1352

32. NCBI Resource Coordinators. Database resources of the National Center for Biotechnology Information. Nucleic Acids Res. 2017;45: D12–D17. doi:10.1093/nar/gkw1071

33. Mashima J, Kodama Y, Fujisawa T, Katayama T, Okuda Y, Kaminuma E, et al. DNA Data Bank of Japan. Nucleic Acids Res. 2017;45: D25–D31. doi:10.1093/nar/gkw1001

34. Alves I, Ambrosini G, Pedone PA, Angelina P, Anisimova M, Appel R, et al. The SIB Swiss Institute of Bioinformatics’ resources: focus on curated databases. Nucleic Acids Res. 2016;44: D27–37. doi:10.1093/nar/gkv1310

35. Burley SK, Berman HM, Kleywegt GJ, Markley JL, Nakamura H, Velankar S. Protein Data Bank (PDB): the single global macromolecular structure archive. In: Wlodawer A, Dauter Z, Jaskolski M, editors. Protein Crystallography. Springer New York; pp. 627–641. doi:10.1007/978-1-4939-7000-1_26

36. Beagrie N, Houghton J. The value and impact of the European Bioinformatics Institute: executive summary. Charles Beagrie Ltd.; 2016.

37. Thessen AE, Patterson DJ. Data issues in the life sciences. Zookeys. 2011; 15–51. doi:10.3897/zookeys.150.1766

38. Costello MJ, Appeltans W, Bailly N, Berendsohn WG, de Jong Y, Edwards M, et al. Strategies for the sustainability of online open-access biodiversity databases. Biol Conserv. 2014;173: 155–165. doi:10.1016/j.biocon.2013.07.042

39. Oliver SG, Lock A, Harris MA, Nurse P, Wood V. Model organism databases: essential resources that need the support of both funders and users. BMC Biol. 2016;14: 49. doi:10.1186/s12915-016-0276-z

40. Kaiser J. Funding for key data resources in jeopardy. Science. 2016;351: 14–14. doi:10.1126/science.351.6268.14

41. Schnoes AM, Brown SD, Dodevski I, Babbitt PC. Annotation error in public databases: misannotation of molecular function in enzyme superfamilies. PLoS Comput Biol. 2009;5: e1000605. doi:10.1371/journal.pcbi.1000605

42. Bengtsson-Palme J, Boulund F, Edström R, Feizi A, Johnning A, Jonsson VA, et al. Strategies to improve usability and preserve accuracy in biological sequence databases. Proteomics. 2016;16: 2454–2460. doi:10.1002/pmic.201600034

43. European Molecular Biology Laboratory, European Bioinformatics Institute. EMBL-EBI Standards and Policies: Third-Party Annotation Policy [Internet]. [cited Jun 28, 2017]. Available: http://www.ebi.ac.uk/ena/about/tpa-policy

44. National Centre for Biotechnology Information. About third-party annotation. In: NCBI [Internet]. [cited Jun 28, 2017]. Available: https://www.ncbi.nlm.nih.gov/genbank/tpa/

45. ten Hoopen P, Amid C, Buttigieg PL, Pafilis E, Bravakos P, Cerdeño-Tárraga AM, et al. Value, but high costs in post-deposition data curation. Database. 2016;2016: bav126. doi:10.1093/database/bav126

46. McQuilton P, Gonzalez-Beltran A, Rocca-Serra P, Thurston M, Lister A, Maguire E, et al. BioSharing: curated and crowd-sourced metadata standards, databases and data policies in the life sciences. Database. 2016;2016. doi:10.1093/database/baw075

47. Jisc. Metadata. In: Jisc [Internet]. 2014 [cited 19 May 2017]. Available: https://www.jisc.ac.uk/guides/metadata

48. Malone J, Stevens R, Jupp S, Hancocks T, Parkinson H, Brooksbank C. Ten simple rules for selecting a bio-ontology. PLoS Comput Biol. 2016;12: e1004743. doi:10.1371/journal.pcbi.1004743

49. Rocca-Serra P, Salek RM, Arita M, Correa E, Dayalan S, Gonzalez-Beltran A, et al. Data standards can boost metabolomics research, and if there is a will, there is a way. Metabolomics. 2016;12: 14. doi:10.1007/s11306-015-0879-3

50. Tenenbaum JD, Sansone S-A, Haendel M. A sea of standards for omics data: sink or swim? J Am Med Inform Assoc. 2014;21: 200–203. doi:10.1136/amiajnl-2013-002066

51. BioSharing [Internet]. 2017 [cited 30 May 2017]. Available: https://biosharing.org

52. Pearson WR, Lipman DJ. Improved tools for biological sequence comparison. Proc Natl Acad Sci U S A. 1988;85: 2444–2448. Available: https://www.ncbi.nlm.nih.gov/pubmed/3162770

53. Cock PJA, Fields CJ, Goto N, Heuer ML, Rice PM. The Sanger FASTQ file format for sequences with quality scores, and the Solexa/Illumina FASTQ variants. Nucleic Acids Res. 2010;38: 1767–1771. doi:10.1093/nar/gkp1137

54. Taylor CF, Field D, Sansone S-A, Aerts J, Apweiler R, Ashburner M, et al. Promoting coherent minimum reporting guidelines for biological and biomedical investigations: the MIBBI project. Nat Biotechnol. 2008;26: 889–896. doi:10.1038/nbt.1411

55. Hinsen K. Platforms for publishing and archiving computer-aided research [version 3; referees: 3 approved]. F1000Res. 2015; doi:10.12688/f1000research.5773.1

56. Piccolo SR, Frampton MB. Tools and techniques for computational reproducibility. Gigascience. 2016;5: 30. doi:10.1186/s13742-016-0135-4

57. Katz D. Is software reproducibility possible and practical? [Internet]. 2017 [cited 19 May 2017]. Available: https://danielskatzblog.wordpress.com/2017/02/07/is-software-reproducibility-possible-and-practical/

58. Hinsen K. Sustainable software and reproducible research: dealing with software collapse [Internet]. 2017 [cited 19 May 2017]. Available: http://blog.khinsen.net/posts/2017/01/13/sustainable-software-and-reproducible-research-dealing-with-software-collapse/

59. Brown CT. How I learned to stop worrying and love the coming archivability crisis in scientific software [Internet]. 2017 [cited 19 May 2017]. Available: http://ivory.idyll.org/blog/2017-pof-software-archivability.html

60. Github [Internet]. 2017 [cited 29 May 2017]. Available: https://github.com/

61. Bitbucket [Internet]. 2017 [cited 29 May 2017]. Available: https://bitbucket.org/

62. Zenodo [Internet]. 2017 [cited 29 May 2017]. Available: https://zenodo.org/

63. Jiménez RC, Kuzak M, Alhamdoosh M, Barker M, Batut B, Borg M, et al. Four simple recommendations to encourage best practices in research software [version 1; referees: 3 approved]. F1000Res. 2017;6. doi:10.12688/f1000research.11407.1

64. Artaza H, Chue Hong N, Corpas M, Corpuz A, Hooft R, Jimenez RC, et al. Top 10 metrics for life science software good practices. F1000Res. 2016;5. doi:10.12688/f1000research.9206.1

65. Wilson G, Bryan J, Cranston K, Kitzes J, Nederbragt L, Teal TK. Good enough practices in scientific computing. PLoS Comput Biol. 2017;13: e1005510. doi:10.1371/journal.pcbi.1005510

66. Faulconbridge A, Burdett T, Brandizi M, Gostev M, Pereira R, Vasant D, et al. Updates to BioSamples database at European Bioinformatics Institute. Nucleic Acids Res. 2014;42: D50–2. doi:10.1093/nar/gkt1081

67. protocols.io [Internet]. 2017 [cited 1 Jun 2017]. Available: https://www.protocols.io/

68. OpenWetWare [Internet]. 2017 [cited 29 May 2017]. Available: http://www.openwetware.org/

69. Schilthuizen M, Vairappan CS, Slade EM, Mann DJ, Miller JA. Specimens as primary data: museums and “open science.” Trends Ecol Evol. 2015;30: 237–238. doi:10.1016/j.tree.2015.03.002

70. Turney S, Cameron ER, Cloutier CA, Buddle CM. Non-repeatable science: assessing the frequency of voucher specimen deposition reveals that most arthropod research cannot be verified. PeerJ. 2015;3: e1168. doi:10.7717/peerj.1168

71. Walters C, Volk GM, Richards CM. Genebanks in the post-genomic age: emerging roles and anticipated uses. Biodiversity. Taylor & Francis; 2008;9: 68–71. doi:10.1080/14888386.2008.9712887

72. Lloyd K, Franklin C, Lutz C, Magnuson T. Reproducibility: use mouse biobanks or lose them. Nature. 2015;522: 151–153. doi:10.1038/522151a

73. Watson PH. Biospecimen complexity - the next challenge for cancer research biobanks? Clin Cancer Res. 2017;23: 894–898. doi:10.1158/1078-0432.CCR-16-1406

74. Schnell S. Ten simple rules for a computational biologist’s laboratory notebook. PLoS Comput Biol. 2015;11: e1004385. doi:10.1371/journal.pcbi.1004385

75. LabTrove [Internet]. 2017. Available: http://www.labtrove.org/

76. Blower JD, Santokhee A, Milsted AJ, Frey JG. BlogMyData: a Virtual Research Environment for collaborative visualization of environmental data. 2010. Available: https://eprints.soton.ac.uk/164533/

77. Walsh E, Cho I. Using Evernote as an electronic lab notebook in a translational science laboratory. J Lab Autom. 2013;18: 229–234. doi:10.1177/2211068212471834

78. Benchling [Internet]. 2017. Available: https://benchling.com/

79. Smith VS, Rycroft SD, Brake I, Scott B, Baker E, Livermore L, et al. Scratchpads 2.0: a Virtual Research Environment supporting scholarly collaboration, communication and data publication in biodiversity science. Zookeys. 2011; 53–70. doi:10.3897/zookeys.150.2193

80. Boettiger C. Migrating to Hugo and Blogdown. In: Boettiger Group Lab Website [Internet]. April 17, 2017 [cited Jun 28, 2017]. Available: http://www.carlboettiger.info/2017/04/19/migrating-to-hugo-and-blogdown/

81. Boettiger C. A reproducible R notebook using Docker. In: Kitzes J, Turek D, Deniz F, editors. The practice of reproducible research: case studies and lessons from the data-intensive sciences. Oakland, CA: University of California Press; 2017. Available: https://www.practicereproducibleresearch.org/case-studies/cboettig.html

82. Project Jupyter. Jupyter Notebook [Internet]. 2014. Available: http://jupyter.org/

83. Rstudio. Rmarkdown [Internet]. 2017. Available: http://rmarkdown.rstudio.com/

84. Docker [Internet]. 2017. Available: https://www.docker.com/

85. Koshland DE Jr. The price of progress. Science. 1988;241: 637. doi:10.1126/science.241.4866.637

86. Jasny BR. Realities of data sharing using the genome wars as case study - an historical perspective and commentary. EPJ Data Science. 2013;2:1. doi:10.1140/epjds13

87. Caetano DS, Aisenberg A. Forgotten treasures: the fate of data in animal behaviour studies. Anim Behav. 2014;98: 1–5. doi:10.1016/j.anbehav.2014.09.025

88. Piwowar HA, Chapman WW. A review of journal policies for sharing research data. Open scholarship: authority, community, and sustainability in the age of Web 20 Proceedings of the 12th International Conference on Electronic Publishing (ELPUB) 2008. Toronto, Canada; 2008. Available: http://ocs.library.utoronto.ca/index.php/Elpub/2008/paper/view/684/0

89. National Research Council, Division on Earth and Life Studies, Board on Life Sciences, Committee on Responsibilities of Authorship in the Biological Sciences. Sharing publication-related data and materials: responsibilities of authorship in the life sciences. National Academies Press; 2003.

90. Oxford University Press. Bioinformatics: Instructions for Authors [Internet]. 2017 [cited 30 May 2017]. Available: https://academic.oup.com/bioinformatics/pages/instructions_for_authors

91. Biomed Central. Editorial policies. In: BioMed Central [Internet]. 2017 [cited 19 May 2017]. Available: http://www.biomedcentral.com/getpublished/editorial-policies

92. Public Library of Science. PLoS Submission Guidelines [Internet]. 2017 [cited 2017]. Available: http://journals.plos.org/plosone/s/submission-guidelines

93. Public Library of Science. PLoS One: Data Availability [Internet]. 2017 [cited 1 Jun 2017]. Available: http://journals.plos.org/plosone/s/data-availability

94. International Committee of Medical Journal Editors. ICMJE: Manuscript Preparation - Preparing for Submission [Internet]. 2017 [cited 1 Jun 2017]. Available: http://www.icmje.org/recommendations/browse/manuscript-preparation/preparing-for-submission.html#d

95. Science. Science: Instructions for Authors [Internet]. 2017 [cited 1 Jun 2017]. Available: http://www.sciencemag.org/authors/instructions-preparing-initial-manuscript

96. Elsevier Inc. Cell Press STAR Methods [Internet]. 2017 [cited 1 Jun 2017]. Available: http://www.cell.com/star-authors-guide

97. Kilkenny C, Browne WJ, Cuthill IC, Emerson M, Altman DG. Improving bioscience research reporting: the ARRIVE guidelines for reporting animal research. PLoS Biology. 2010;8: e1000412. doi:10.1371/journal.pbio.1000412

98. Naughton L, Kernohan D. Making sense of journal research data policies. Insights. UKSG in association with Ubiquity Press; 2016;29. doi:10.1629/uksg.284

99. Iqbal SA, Wallach JD, Khoury MJ, Schully SD, Ioannidis JPA. Reproducible research practices and transparency across the biomedical literature. PLoS Biol. 2016;14: e1002333. doi:10.1371/journal.pbio.1002333

100. Nekrutenko A, Taylor J. Next-generation sequencing data interpretation: enhancing reproducibility and accessibility. Nat Rev Genet. 2012;13: 667–672. doi:10.1038/nrg3305

101. Ioannidis JPA, Khoury MJ. Improving validation practices in “omics” research. Science. 2011;334: 1230–1232. doi:10.1126/science.1211811

102. Errington TM, Iorns E, Gunn W, Tan FE, Lomax J, Nosek BA. An open investigation of the reproducibility of cancer biology research. Elife. 2014;3. doi:10.7554/eLife.04333

103. Nature Publishing Group. Scientific Data: Information for Authors [Internet]. 2017 [cited 2 Jun 2017]. Available: https://www.nature.com/sdata/publish/for-authors

104. Oxford University Press. GigaScience: Data Note [Internet]. 2017 [cited 2 Jun 2017]. Available: https://academic.oup.com/gigascience/pages/data_note

105. Wolpert AJ. For the sake of inquiry and knowledge—the inevitability of open access. N Engl J Med. Mass Medical Soc; 2013;368: 785–787. Available: http://www.nejm.org/doi/full/10.1056/NEJMp1211410

106. Laakso M, Welling P, Bukvova H, Nyman L, Björk B-C, Hedlund T. The development of open access journal publishing from 1993 to 2009. PLoS One. 2011;6: e20961. doi:10.1371/journal.pone.0020961

107. DCMI Usage Board. DCMI Metadata Terms. In: Dublin Core Metadata Initiative [Internet]. 14 Jun 2012 [cited 2 Jul 2017]. Available: http://dublincore.org/documents/dcmi-terms/

108. ORCID [Internet]. 2017 [cited 2 Jun 2017]. Available: https://orcid.org/

109. HUGO Gene Nomenclature Committee. Journals [Internet]. 2017 [cited 2 Jun 2017]. Available: http://www.genenames.org/useful/journals

110. McMurry JA, Juty N, Blomberg N, Burdett T, Conlin T, Conte N, et al. Identifiers for the 21st century: how to design, provision, and reuse persistent identifiers to maximize utility and impact of life science data. PLoS Biol. 2017;15: e2001414. doi:10.1371/journal.pbio.2001414

111. Genetics Society of America. Why publish in the GSA journals? [Internet]. 2017 [cited 2 Jun 2017]. Available: http://www.genetics.org/content/why

112. Australian National Data Service. ANDS Guide: De-identification [Internet]. 2017. Available: http://www.ands.org.au/__data/assets/pdf_file/0003/737211/De-identification.pdf

113. Australian Government, National Health and Medical Research Council. National Statement on ethical conduct in human research [Internet]. 2007 - updated 2015. Available: https://www.nhmrc.gov.au/_files_nhmrc/publications/attachments/e72_national_statement_may_2015_150514_a.pdf

114. Australian National Medical Research Data Storage Facility. Legal, best practice and security frameworks for consideration in operation of the Australian National Medical Research Data Storage Facility [Internet]. 2016 Jan. Available: http://med.data.edu.au/wp-content/uploads/2016/01/Med.Data_Legal_Security_Framework_Jan2016.pdf

115. Baker M, Keeton K, Martin S. Why traditional storage systems don’t help us save stuff forever. Proc 1st IEEE Workshop on Hot Topics in System Dependability. 2005. pp. 2005–2120. Available: http://www.hpl.hp.com/techreports/2005/HPL-2005-120.pdf?jumpid=reg_R1002_USEN

116. European Bioinformatics Institute. EMBL-EBI Train online. In: EMBL-EBI Train online [Internet]. [cited 19 May 2017]. Available: https://www.ebi.ac.uk/training/online/

